# The Atlas of Inflammation-Resolution (AIR)

**DOI:** 10.1101/2020.01.27.921882

**Authors:** Charles N. Serhan, Shailendra Gupta, Mauro Perretti, Catherine Godson, Eoin Brennan, Yongsheng Li, Oliver Soehnlein, Takao Shimizu, Oliver Werz, Valerio Chiurchiù, Angelo Azzi, Marc Dubourdeau, Suchi Smita Gupta, Patrick Schopohl, Matti Hoch, Dragana Gjorgevikj, Faiz M. Khan, David Brauer, Anurag Tripathi, Konstantin Cesnulevicius, David Lescheid, Myron Schultz, Dirk Repsilber, Robert Kruse, Angelo Sala, Jesper Z. Haeggström, Bruce D. Levy, János G. Filep, Olaf Wolkenhauer

## Abstract

Acute inflammation is a protective reaction by the immune system in response to invading pathogens or tissue damage. Ideally, the response should be localized, self-limited, and returning to homeostasis. If not resolved, acute inflammation can result in organ pathologies leading to chronic inflammatory phenotypes. Acute inflammation and inflammation resolution are complex coordinated processes, involving a number of cell types, interacting in space and time. The biomolecular complexity and the fact that several biomedical fields are involved, make a multi and interdisciplinary approach necessary.

This Atlas of Inflammation Resolution (AIR) is a web-based resource capturing the state-of-the-art in acute inflammation and inflammation resolution research. The AIR provides an interface for users to search thousands of interactions, arranged in inter-connected multi-layers of process diagrams, covering a wide range of clinically relevant phenotypes. The AIR serves as an open access knowledgebase, including a gateway to numerous public databases. It is furthermore possible for the user to map experimental data onto the molecular interaction maps of the AIR, providing the basis for bioinformatics analyses and systems biology approaches. By mapping experimental data onto the Atlas, it can be used to elucidate drug action as well as molecular mechanisms underlying different disease phenotypes. For the visualization and exploration of information, the AIR uses the Minerva platform, which is a well-established tool for the presentation of disease maps. The molecular details of the AIR are encoded using international standards.

The Atlas of Inflammation Resolution was created as a freely accessible resource, supporting research and education in the fields of acute inflammation and inflammation resolution. The AIR connects research communities, facilitates clinical decision making, and supports research scientists in the formulation and validation of hypotheses.

## 1. Introduction

### 1.1 Problem and research gap

The acute inflammatory response is the first protective reaction mounted by the host tissue against invading pathogens, foreign bodies and/or injury^1^. Acute inflammation is a highly coordinated, active, nonlinear spatial-temporal process for the removal of invading pathogens and the repair of damaged tissues to reestablish homeostasis^2–4^. If the acute inflammatory response is not resolved, it can contribute to organ pathology and amplify many widely occurring chronic inflammatory clinical phenotypes including arthritis, neurodegenerative diseases, metabolic syndrome, asthma, allergy, diabetes, aging, cancers, organ fibrosis, cardiovascular and periodontal diseases^5,6,15,7–14^.

The landscape of the entire acute inflammatory response can be broadly divided into four phases, namely, i) inflammation initiation, ii) transition, iii) resolution and iv) return to a new state of homeostasis. The boundaries of all these phases are appreciated at the cellular and tissue level and are just being defined at the molecular and biomarker levels. The physiological terrain leading to acute inflammatory responses contains a large number of pro-inflammatory mediators (PIM) and specialized pro-resolving mediators including specialized pro-resolving lipid mediators (SPM), proteins, peptides, autocoids, varieties of innate immune cells and a vast number of regulators (molecular switches) in the form of feedback and feedforward loops, making the whole system dynamic and complex (Figure 1).

**Figure 1:**
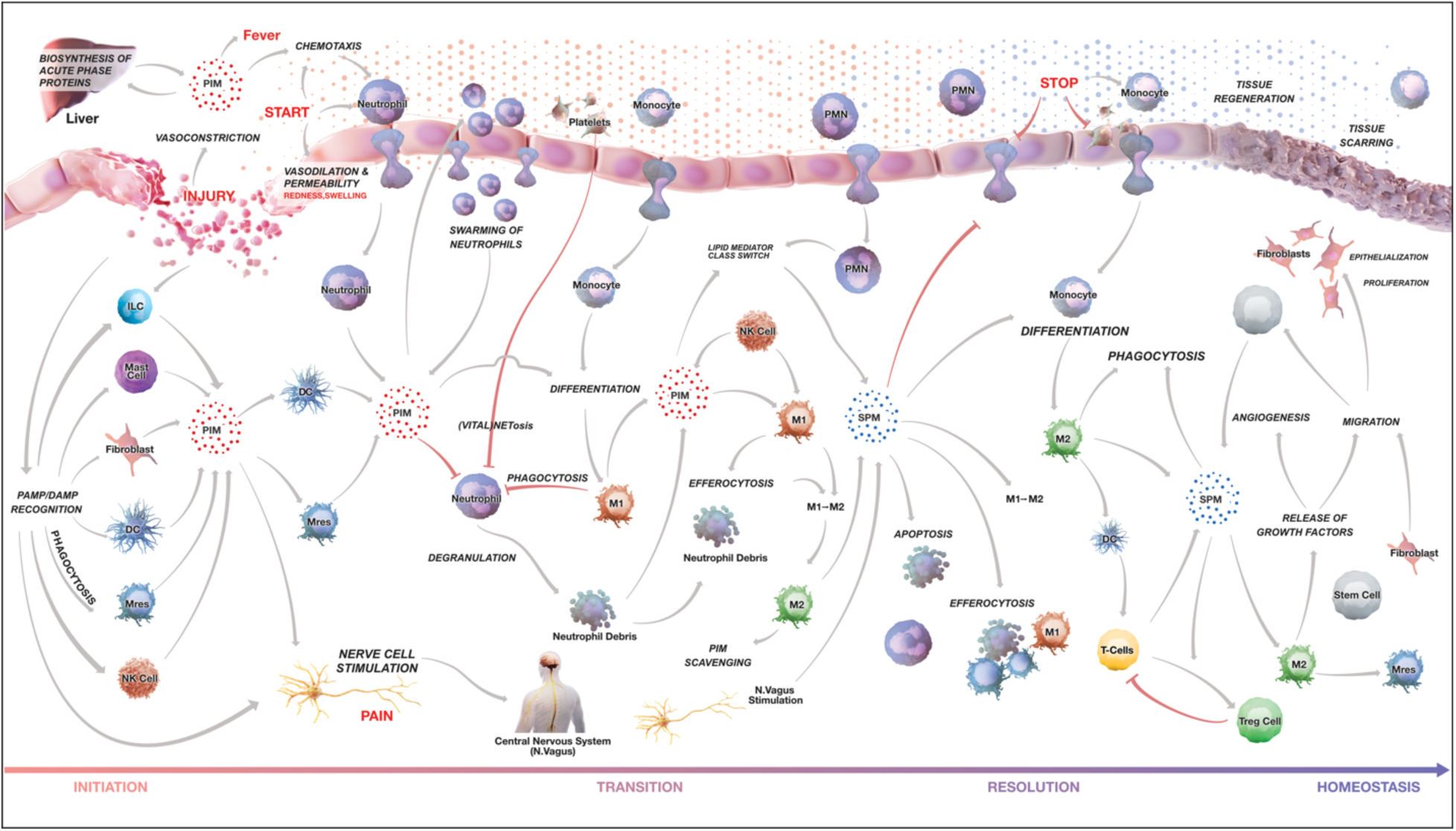
Phenotype level representation of the AIR. The landscape of acute inflammatory response is divided into four overlapping phases: Initiation, transition, resolution and return to tissue homeostasis. Interactions between immune cell types, vascular endothelial cells, mucosal epithelial cells as well as the associated processes and phenotype are depicted. Arrows indicate information flow. The color of arrows indicates regulation type; gray for activation and red for inhibition. Each process is connected with underlying manually curated and annotated molecular interaction maps (see text).

Understanding the interplay between different regulators is necessary to understand immune cell function and phenotypic adaptations during acute inflammation and its resolution. Perturbation of these sensitive chemical mediator networks and molecular switches, e.g. through mutations or dysregulated expression, can lead to a misrouted behavior to clear or eradicate the initiating stimulus and ultimately cause the emergence of chronic diseases^9,16–18^.

To conceptually analyze and intuitively visualize interactions in such a vast regulatory network, the construction of a molecular interaction map is the first step in modern biological studies^19,20^. A number of comprehensive regulatory maps have been constructed to help in the understanding of the mechanisms underlying complex biological systems^21–24^, however, only a handful of attempts have been made so far to develop a mechanistic understanding of complex inflammatory disorders through the investigation of underlying molecular interaction maps^25–28^.

Research communities working on inflammation and inflammation-resolution are highly diverse typically focusing on either one immune cell type, a specific mediator, or pathway associated with any specific disease pathophysiology under investigation. Intriguingly, the presence of nonlinear interactions between immune cells and associated pathways renders the prediction of outcomes highly non-intuitive and requires a bird’s-eye view of the whole system. Moreover, to this date, there is no common web platform wherein a community can share their viewpoints on the impact that acute inflammatory phenotypes may have on the emergence of specific disease settings. It would be prudent and apt to design a multi-layered representation of inflammation and inflammation resolution, wherein manipulation of one or more parameters in one stratum would automatically predict the resulting mechanistic changes in another layer.

### 1.2 Development and target groups

We conceptualized the Atlas of Inflammation Resolution (AIR) as a community resource to connect clinicians, scientists, students, and pharmaceutical companies working in the area of acute and chronic inflammation. This can be realized only when the AIR is able to support 1) clinicians in decision making (e.g. patient stratification; predicting response to therapy; therapy personalization; prognosis prejudgment), 2) research scientists in the development of hypotheses for experimental design (e.g. identification of molecular switches, mapping of high-throughput experimental data, understanding mechanisms), and 3) pharmaceutical companies in the identification of new therapeutic targets. The multi-level nature of the systems interacting as presented in the AIR can serve as a starting point for defining and designing acute inflammation and inflammation resolution related dynamic mathematical models. The AIR would naturally be useful for education and training purposes as well. With this in mind, we present the AIR as a novel community resource.

### 1.3 Technology and usages

AIR allows users to visualize detailed molecular level events underlying biological processes, pathways and cell/tissue-related phenotypes associated with acute inflammation initiation, transition, resolution and finally return to homeostasis. All the submaps are prepared using standard Systems Biology Markup Language (SBML)^29^ to ensure their reusability and are accessible through the MINERVA platform^30^ using a web-browser. The AIR is a portal which enables connection to state-of-the-art publicly available databases, providing information about genes, proteins, lipids, miRNAs, lncRNAs, chemicals and drugs. It is furthermore a tool to highlight research gaps and supports the formulation of new hypotheses to address those gaps.

### 1.4 Content of the manuscript

In the following sections, we describe the methods and workflow used for the construction of AIR along with the functionality that users will be provided with when accessing the AIR. The paper also details the use of AIR by clinicians, scientists, students, and pharmaceutical industries.

## 2. Methods

### 2.1 Construction of AIR combining bottom-up and top-down approaches

Classical disease networks are designed using a bottom-up approach, where the phenotype is represented by interacting subsystems (functional modules), each containing evidence-based molecular interactions of clinical relevance^31^. With the availability of omics data, disease networks are frequently built top-down, where signatures (e.g. differentially expressed genes) are being first identified and subsequently on connected to their interacting and regulatory partners. All the connected components are then analyzed to access overrepresented molecular processes and pathways, which together are supposed to coordinate for the emergence of disease phenotype^32,33^. Because of the pros and cons associated with both approaches, the AIR is conceptualized as a “middle-out” approach where the best bottom-up model is integrated with the largest top-down model^31^ (Figure 2).

**Figure 2:**
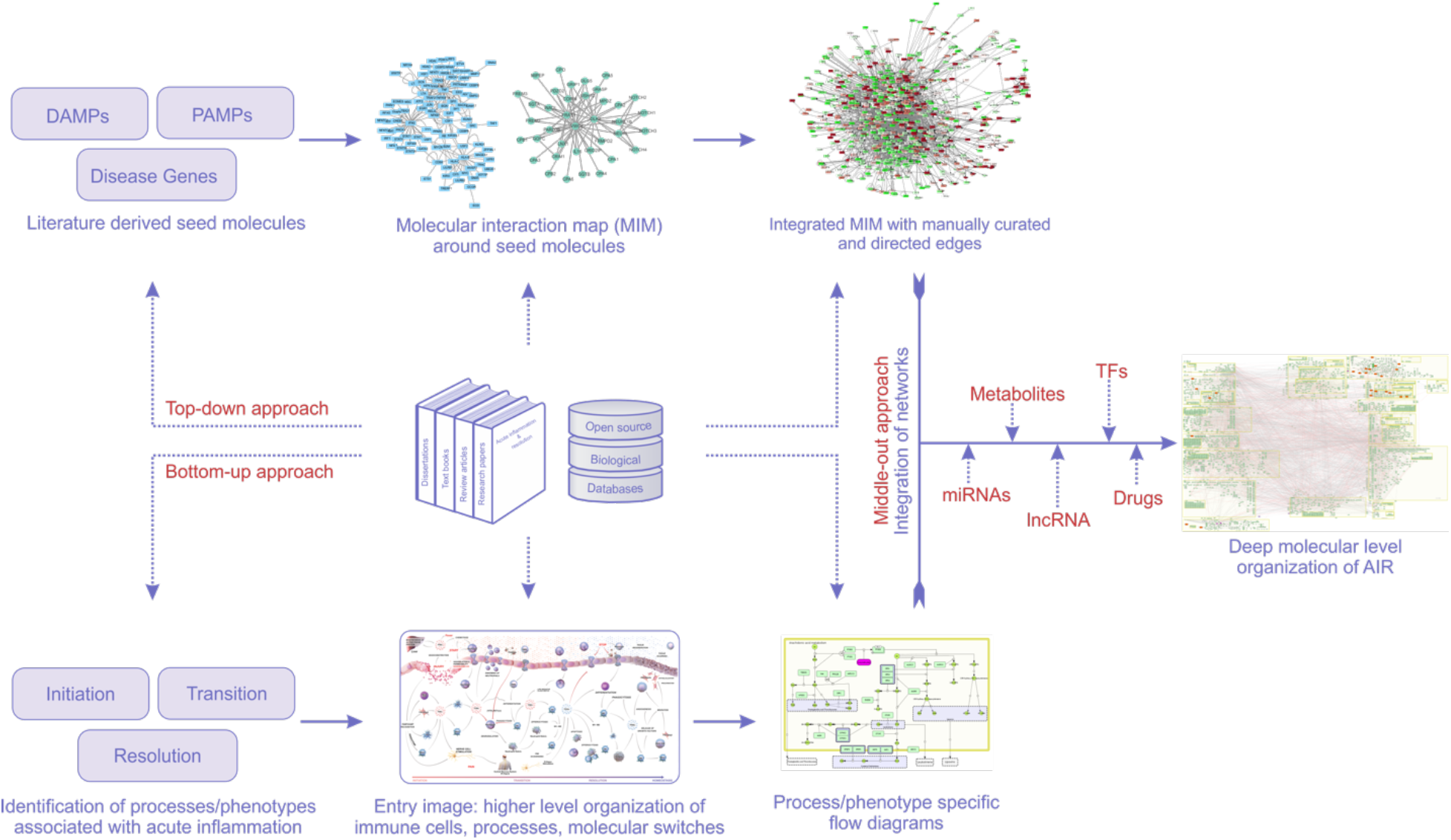
Workflow for the construction of the Atlas of Inflammation Resolution (AIR). The AIR is constructed both bottom-up and top-down. In case of the bottom-up approach, higher level processes, phenotypes and interplay between immune cells were identified in various stages of acute inflammation. These processes and phenotypes were extended in the form of information flow diagrams in standard SBML notations. In the top-down approach, first seed molecules were identified from damage-associated molecular patterns (DAMPs), Pathogen-associated molecular patterns (PAMPs) and key disease genes associated with selected clinical phenotypes of acute inflammation. Each seed molecule is then extended with the experimentally validated interacting partners. Models generated using bottom-up and top-down approaches were later merged together as a middle-out approach to create a comprehensive molecular interaction layer of AIR. This layer is further extended with experimentally validated regulatory layers of transcription factors, miRNAs, lncRNAs, drugs and metabolites.

In case of the bottom-up approach, we first reviewed published literature to identify molecular processes, cell and tissue specific phenotypes associated with one of the four acute inflammatory phases i.e. inflammation initiation, transition, resolution and return to homeostasis (Figure 1). For each of the processes/phenotypes, we manually screened literature and databases (Reactome (https://reactome.org), KEGG (https://www.genome.jp/kegg/), InnateDB (https://innatedb.com)) to extract experimentally validated signaling and regulatory events. These are finally represented by a process diagram using the CellDesigner software (http://www.celldesigner.org) in standard systems biology graphics notation (SBGN) representation.

The top-down approach to the acute inflammation response started with the collection of key molecular signatures (seed molecules). These molecular signatures are then extended with known interacting and regulatory partners. As acute inflammation can be triggered by a variety of etiological agents, from which we considered three sets of seed molecules for the construction of AIR, these are 1) damage associated molecular patterns (DAMPs); 2) receptors recognizing pathogen associated molecular patterns (PAMPs); and 3) disease genes from selected acute inflammatory clinical phenotypes. While DAMPS and PAMPs-recognizing receptors were mainly identified through research articles, key molecules associated with acute inflammatory clinical indications were screened through disease-gene association databases. To this end, we mainly used DisGeNET (https://disgenet.org), eDGAR (http://edgar.biocomp.unibo.it), KEGG disease (https://www.genome.jp/kegg/disease) databases. As many of the disease-gene association databases are based on text mining algorithms, we manually cross checked the disease-gene association from associated research papers. For each of the seed molecules, we identified interacting molecular components from literature and databases to prepare a molecular interaction network. We also used the Bisogenet 3.0.0 app^34^ available on Cytoscape 3.7.0 which connects large number of biological databases (e.g. DIP (https://dip.doe-mbi.ucla.edu/), BioGRID (https://thebiogrid.org), HPRD (https://hprd.org), IntAct (https://www.ebi.ac.uk/intact/), MINT (https://mint.bio.uniroma2.it)) to first create biological networks around each of the seed molecule. We extracted only experimentally validated interactions.

Networks generated using bottom-up and top-down approaches were merged together as a middle-out approach to present a comprehensive molecular interaction network of acute inflammation on AIR. This middle-out approach not only enabled us to expand various acute inflammatory processes/phenotypes (identified in bottom-up approach) with underlying molecular level interactions but also helped in annotating several molecular interactions (top-up approach) with specific process/phenotype.

In addition, many chemical mediators play central roles in the onset of inflammation and later in its resolution. Considering this, we have currently included biosynthesis pathways and downstream signaling cascades of PIM and SPM into the AIR.

Experimentally validated regulatory layers, which include miRNAs from miRbase (http://www.mirbase.org), miRTarBase (http://mirtarbase.mbc.nctu.edu.tw), TriplexRNA (https://triplexrna.org); transcription factors from TRNSFAC (http://genexplain.com/transfac), TRRUST (https://www.grnpedia.org/trrust) and HTRIdb (http://www.lbbc.ibb.unesp.br/htri); long noncoding RNAs from EVLncRNAs (http://biophy.dzu.edu.cn/EVLncRNAs), lncRNADisease (http://www.cuilab.cn/lncrnadisease) databases are also integrated with the AIR using an inhouse script. The overall workflow for the construction of AIR is described in Figure 2.

### 2.2 AIR as a directed graph

Providing direction (e.g. activation, inhibition) to the edges connecting various nodes in the network is a crucial step for network topological analyses and for initiating dynamic systems biological models for the prediction of biomarkers and therapeutic candidates. Only a directed graph will provide mechanistic insights through the study of network motifs and system dynamics. Giving direction to interactions is however a manual effort. Databases using text mining approaches are prone to include false positive information about regulatory directions. We therefore manually cross-checked associated publications before providing directions to the connected edges. More than 80% of the total interactions in the network are directed as of December 2019. The AIR is not just an addition to the existing knowledgebase around acute inflammation but also partially parameterized which can be easily customized for context-specific (e.g. species, cell types, genetics, environment) dynamical model to understand detailed insights of acute inflammation and resolution. While such systems biology approaches are already well established in cancer research^23^, acute inflammation and inflammation resolution offer plenty new opportunities for more interdisciplinary approaches using mathematical modelling and computer simulations. The AIR provides a valuable starting point to identify core regulatory networks that can be subjected to dynamic mechanistic modelling. The entire molecular interaction map of the AIR is encoded using standardized representations, allowing the use of bioinformatics approaches to study it, using graph theoretic and statistical approaches. The Minerva platform allows the integration of plugins for this purpose.

### 2.3 Annotation of AIR content

We enriched every gene/protein in the AIR with specific UniProt-, HGNC-, RefSeq-, Ensembl- and NCBI-ID as well as common aliases with the full name of the encoding protein. In case of small molecules, the ChEBI-ID is provided. Every interaction in AIR is hyperlinked to the respective literature and database. With all the manual curation and annotation, we make AIR a reliable resource for studying processes related to acute inflammation and its resolution.

### 2.4 AIR technical implementation

We used OpenLayer and Google Maps based techniques to bring the AIR to the web-browser for easy visualization implemented on the MINERVA platform developed by University of Luxembourg ^35^. Considering various user groups (clinicians, research scientists, pharmaceutical companies), we divided the AIR into three layers (Figure 3). The <u>top</u> layer consists of immune cell types, cellular processes and compartments. <u>Middle or process layer</u> is comprised of submodules and biomolecular species and finally the <u>bottom layer</u> provides information about molecular level interactions. To reduce the computational effort in processing network images and the requirements to the high-speed local internet, we tiled the CellDesigner representation of the AIR. Tiling is a technique to cut images into a matrix of smaller images of the same size and store it in a specific folder structure. With this, only the currently viewed parts of the AIR can be loaded and displayed. We wrote small Python scripts for tiling and layering to test if the CellDesigner export of the map fits to the postproduction process of the AIR. A local MINERVA instance was installed and tested with AIR for various security and reliability issues.

**Figure 3:**
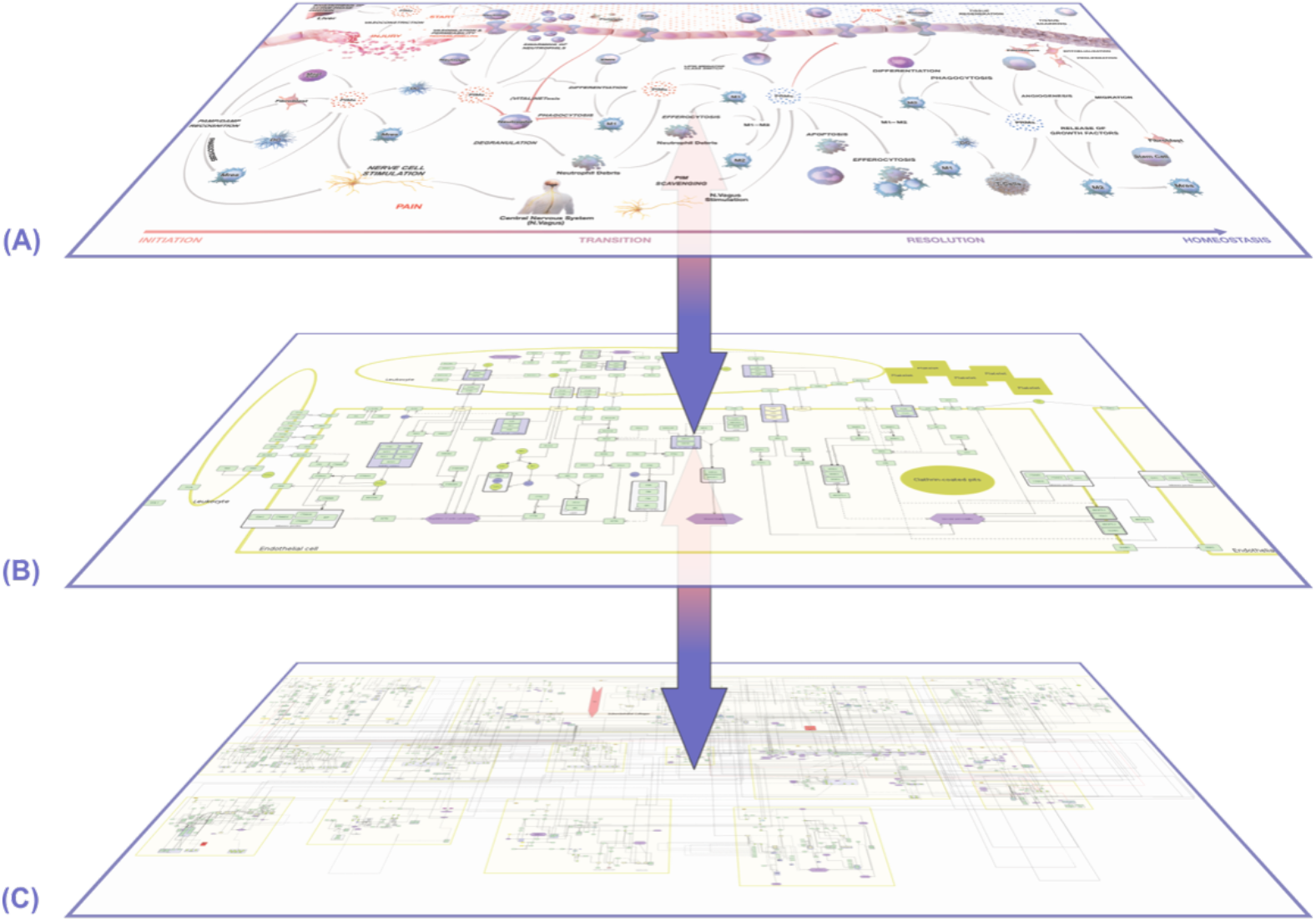
Hierarchical organization of AIR. (A) The top phenotype layer contains immune cell types, cellular processes/phenotypes and tissue level organization. Clinicians are generally interested in connecting their patient data to this layer. (B) Each process in the top layer is connected to a respective signal flow diagram. The process layer describes key molecules/pathways regulating top layer. This layer is suitable for research scientists to generate new hypotheses on the mechanistic insights of disease phenotype regulation. (C) The lower layer contains a comprehensive Molecular Interaction Map (MIM) where all the processes are merged together at the molecular level. The layer is also enriched with currently available experimentally validated regulatory information. Each layer provides an opportunity to map and analyze specific data (e.g. Top layer: FACS analysis; middle layer: immune signaling; bottom layer: multiomics data). Due to the communication across multiple layers, the AIR provides a platform for integrative data analysis.

**Figure 3:**
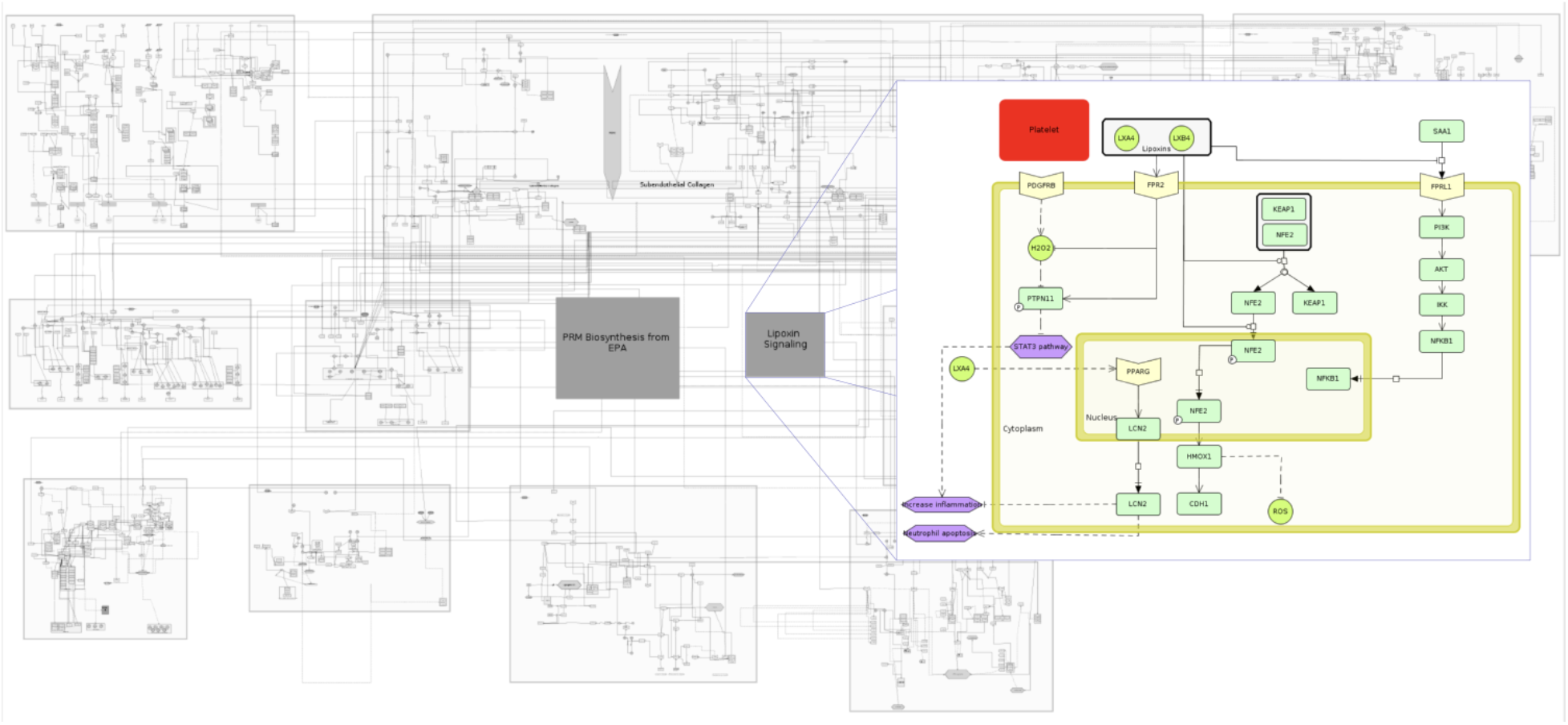
The deepest molecular level layer of the AIR. The layer indicates that all the submaps present in the AIR are highly connected. Various submaps are also integrated together with transcription factors, miRNAs, lncRNAs, chemicals and drugs molecules which are represented as inter submaps species. A detailed molecular interaction map of Lipoxin Signaling in platelets is shown for reference.

## 3. Results

### 3.1 Entry to the AIR

The AIR is a collection of molecular, subcellular, cellular, tissue and functional maps in the context of acute inflammation and inflammation resolution. Depending on the user’s need, the AIR provides various entry points to access all the curated information. AIR is designed with the aim to serve diverse communities including clinicians, research scientists and drug developers from pharmaceutical industry. Clinicians generally rely on the higher-level organization, i.e. at the cellular and tissue level to understand the pathophysiology of disease. They are interested in either classifying patients into various risk groups or in therapy scheduling based on patient’s pathological reports. Time-series transcriptomics profiling of patients is getting popular day-by-day in hospitals which motivate clinicians to use the data in therapy personalization without involving themselves in a data analysis pipeline ^36^. On the other hand, investigators are interested in generating new hypotheses for the design of experiments related to the phenotype under observation. AIR provides multiple layers to extract all of this multi-level information as shown in Figure 3.

### 3.2 Description of the various submaps available on AIR

For better navigation and visualization, the AIR is divided into four phases; these are: i) inflammation initiation; ii) transition; iii) resolution and iv) return to a new state of function and homeostasis.

The phase of **inflammation initiation** starts with the onset of acute inflammation (e.g. invasion by pathogens, tissue damage due to injury or operation etc.), operated by the innate branch of immunity, where the body recognizes DAMPs and PAMPs and releases various PIM and acute phase proteins (APPs) to initiate immune responses. The detailed molecular level events are summarized in submaps ‘DAMPs and PAMPs recognition’ and ‘Regulation of APPs’. Selected chemoattractants and APPs trigger the production of various PIM such as chemokines and others, which in turn are responsible for immediate vasoconstriction followed by vasodilation and increased vasopermeability to facilitate the recruitment of neutrophils to fight against invading pathogens. Various innate immune cells, such as tissue resident macrophages (M_res_), dendritic cells (DC), mast cells and fibroblasts also trigger the production of PIM. In the AIR, molecular level events summarizing these processes are shown in the submaps ‘Chemotaxis’, ‘Vasoconstriction, vasodilation and permeability’, ‘Biosynthesis of PIM’, and ‘Leukocyte adhesion and transmigration’.

Acute inflammation is commonly recognized by the five cardinal signs (Redness, Heat, Swelling, Pain and impaired function) known to ancient physicians^1,37^. These cardinal signs are also integrated in various submaps. For the **inflammation transition** phases, neutrophil swarming was considered as an active process where large number of neutrophils makes a network around a site of acute inflammation. As detailed molecular mechanisms underlying neutrophil swarming are not known, we provide a separate submap describing this process. The aim here is to strengthen all the submaps and connecting missing links with the help of broad inflammation and inflammation-resolution communities. The myriad of cellular processes associated with neutrophils and other white blood cells, such as neutrophil apoptosis, phagocytosis, efferocytosis etc. are also shown in subsequent submaps.

The AIR describes two important switches (macrophage polarization to the broadly defined functionally distinct cell types transitions (e.g. M1 to M2) and lipid mediator class switch) as detailed submaps which are mainly responsible for initiating the **inflammation resolution** phase. Specifically, for the lipid mediator class switch, the AIR provides detailed molecular events and reactions associated with the production of PIM and SPM from arachidonic acid (AA), eicosapentaenoic acid (EPA) and docosahexaenoic acid (DHA)^18,38^ in separate submaps. Other submaps which are associated with inflammation resolution phase are ‘STOP signals for platelet aggregation’ links to the coagulation cascade^39^, ‘STOP signal for neutrophil adhesion’, ‘Monocyte differentiation’ and ‘T_h_-cell signaling cascade, linking resolution to adaptive immunity’.

For the phase of **homeostasis and return to tissue function**, we included processes, such as ‘Tissue scaring and regeneration’, ‘Epithelialization’, ‘Stem cell recruitment and proliferation’ and ‘Angiogenesis’ in subsequent submaps. Submaps currently available on the AIR are listed in Table 1.

**Table 1:** List of key submaps available in the AIR

| Phase | Submap title |
| --- | --- |
| Initiation | Pathogen associated molecular pattern (PAMPs) signaling |
|  | Vasodilation vasoconstriction and permeability |
|  | Leukocyte adhesion and transmigration |
|  | Neutrophil chemotaxis |
|  | Natural killer cell chemotaxis |
|  | Biosynthesis of prostaglandins, thromboxanes and leukotrienes from arachidonic acid |
|  | Biosynthesis of prostaglandins from EPA |
| Transition | Neutrophil apoptosis |
|  | Macrophage phagocytosis |
|  | Monocyte transmigration |
|  | Efferocytosis |
| Resolution | Macrophage M1 to M2 class switch |
|  | Biosynthesis of lipoxins from arachidonic acid |
|  | Biosynthesis of resolvins, maresins, protectins and their conjugates from DHA |
|  | Biosynthesis of E-class resolvins and lipoxins from EPA |
|  | Resolvins mediated signaling cascade |
|  | Protectins mediated signaling cascade |
|  | Maresins mediated signaling cascade |
|  | Lipoxins mediated signaling cascade |
|  | STOP signaling |
| Homeostasis and return to tissue function | Wound healing |

In all submaps, genes and proteins are described by official symbols provided by HUGO gene nomenclature committee (HGNC). All metabolites are provided with ChEBI official names. This helps in connecting all the submaps together as a base layer as shown in Figure 3.

### 3.3 Acute inflammation and its resolution as a coordinated process

The creation of the AIR demonstrates that inflammation resolution is a multilevel spatio-temporal process. This view suggests the application of approaches from systems theory to the field of inflammation resolution. As our body responds to the perturbations initiated by damage, injury to the tissues or invasion by pathogens, the main goal is to regain homeostasis through the coordination of several subprocesses each of which is regulated by large numbers of immune cells and molecules. Underlying the notion of ‘regulation’ is the existence of feedback loops, which in the AIR are found through loops in the directed graph that is the lower molecular interaction map layer. The imbalance in the performance of each subprocess in the lower layer is balanced by the coordination layers which provide signals across subprocesses. These coordination layers in acute inflammation resolution can be summarized as the molecular switches, such as lipid mediator class switch which are responsible for the production of either PIM or SPM from the same precursor. The concept of Interaction Balance Coordination Principle, originally developed by Mesarovic and colleagues^40,41^ serves here as a conceptual model for multilevel coordination (Figure 4). The AIR provides a platform to investigate the dynamics of regulatory subprocesses and coordination layer events in the context of different clinical disease phenotypes.

**Figure 4:**
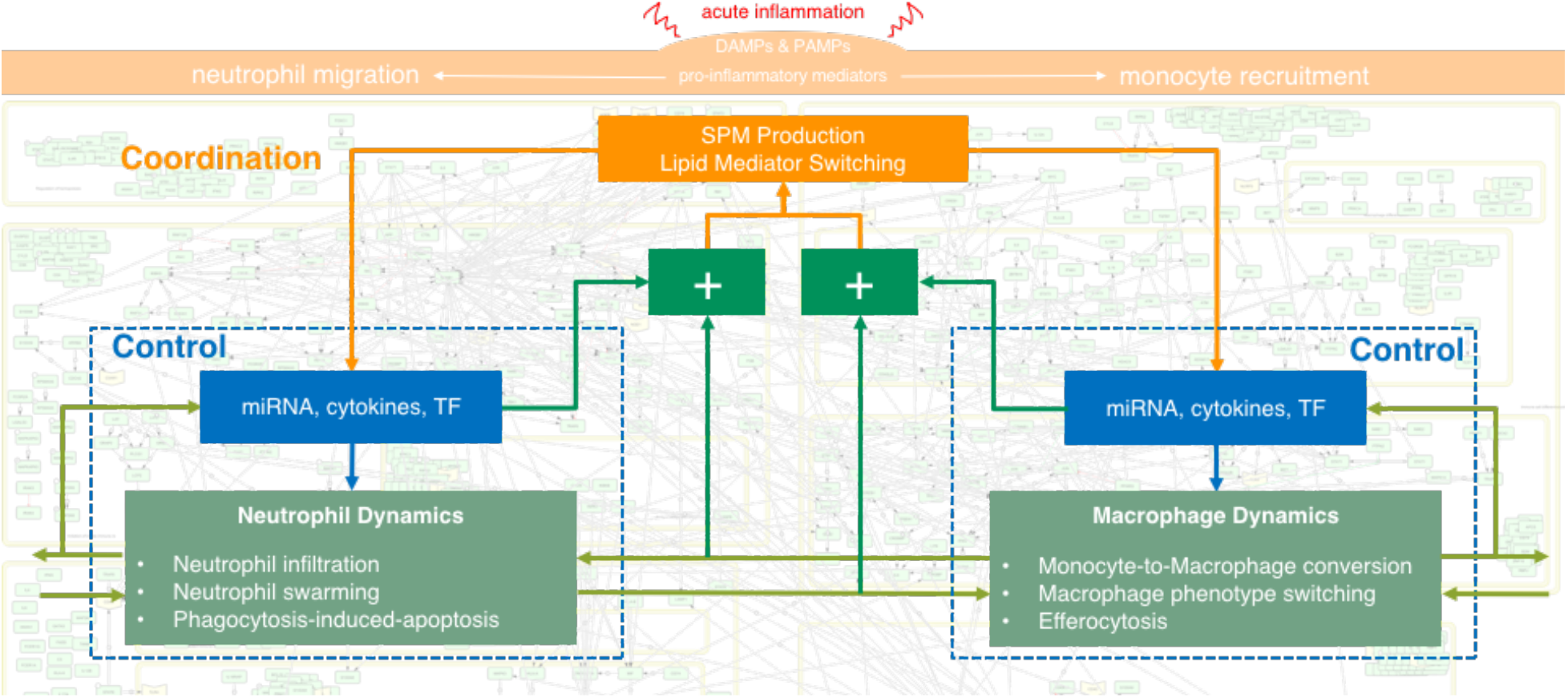
Acute inflammation and inflammation resolution follows the concept of Mesarovic’s Interaction Balance Coordination Principle. Two subprocesses (neutrophil dynamics and macrophage dynamics) are controlled by their respective miRNA, cytokines and transcription factors (TF). These subprocesses communicate together in the regulation of phenotype. If there is any imbalance in the desired and actual outputs by these processes (shown by ‘+’ sign, higher level coordination layer (shown here by ‘SPM Production’, ‘Lipid Mediator Switching’) provide signals and make the balance between subprocesses to return to homeostasis. The AIR provides molecular level details of these coordination layers and offers an opportunity to harness these layers for therapeutic purposes.

### 3.4 Potential uses of the AIR

#### 3.4.1 The AIR as a portal to connect public databases

All biomolecules and chemicals present in the AIR are manually annotated with official names (official HGNC symbol for genes and proteins, ChEBI name for drug and chemicals) and reactions are manually annotated with PubMed IDs. The majority of the state-of-art databases can be linked out directly by selecting nodes or reactions present in the AIR (Figure 5).

**Figure 5:**
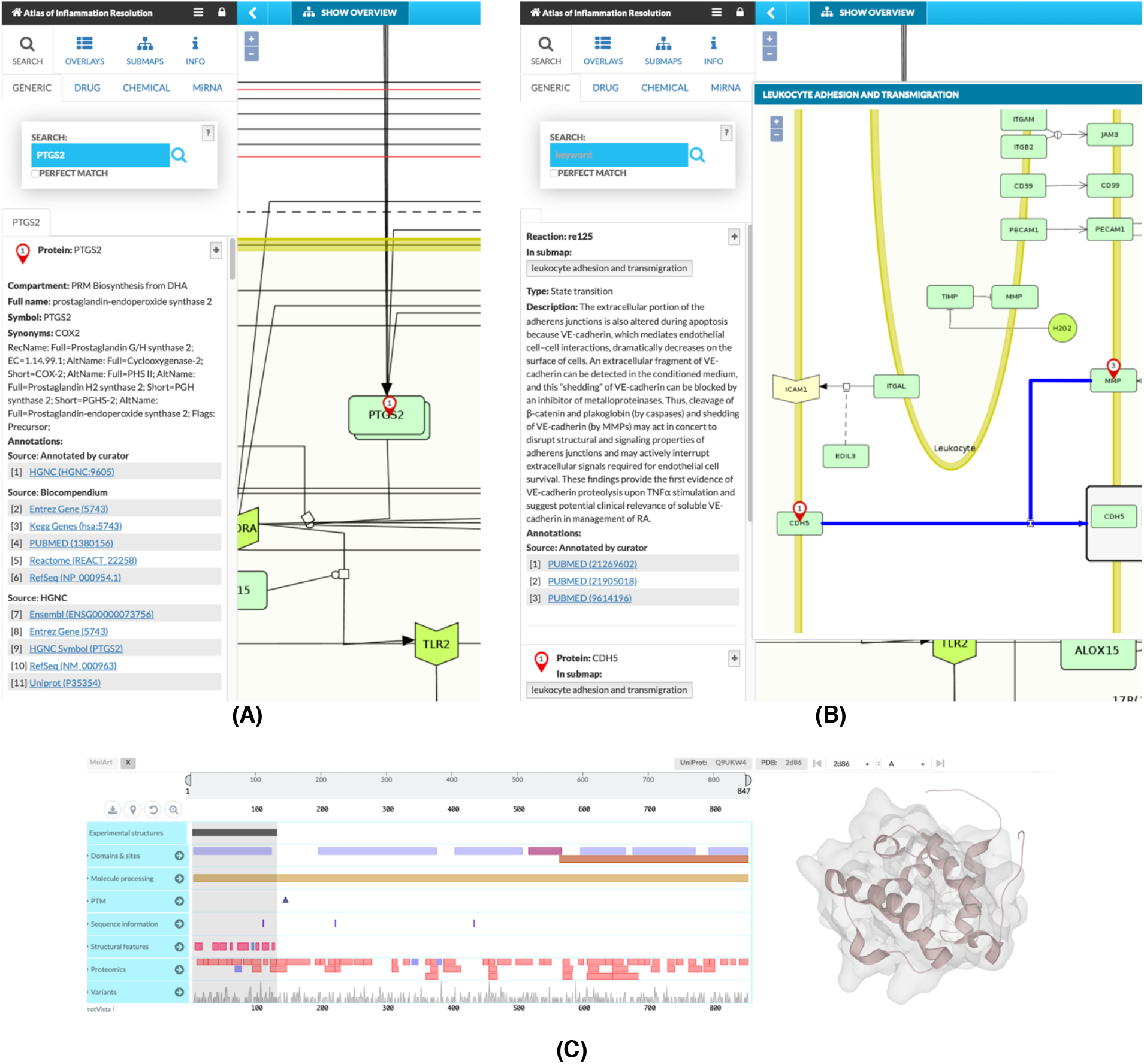
The AIR as a portal to connect public databases. A) A node ‘PTGS2’ is selected. All the link outs to various state-of-the-art databases (HGNC, Entrez gene, KEGG gene, Pubmed, Reactome, Refseq, Uniprot etc.) are directly available on the left panel. (B) A reaction is selected and the literature from where the reaction is derived is provided as Pubmed IDs. Wherever possible, we also summarize the reaction in the context of acute inflammation and inflammation resolution in the description section. (C) Snapshot of MolArt plugin integrated with MINERVA interface. 3D structure of the CH domain of VAV-3 protein is shown as an example.

#### 3.4.2 The AIR as a ready-to-use resource from structure analysis to dynamic models

All the submaps available on the AIR are prepared in standard SBML notation along with complete annotation using CellDesigner tools to ensure their reusability. These submaps can be directly downloaded from the AIR and used for *in silico* simulation, perturbation experiments or network analysis. The majority of the proteins that have a 3D structure already resolved and available in Protein Data Bank can be directly visualized using MolArt visualization plugin^42^ integrated in MINERVA environment (Figure 5C). This plugin helps users in exploring both sequence and structural features (including protein variation data from large scale studies) associated with proteins. In addition to the sequence and structure level features of the protein, one can connect 3D structures of protein complexes, drugs and chemicals bound to their receptor proteins which can be directly integrated into structure-based drug development pipelines. Users can develop their own Minerva plugins to analyze the content of the MIM.

#### 3.4.3 Visualization of time-series omics data

AIR hosted on the MINERVA platform provides the user with an interactive interface to map time-series omics data. Once uploaded on the AIR, these data can be visualized on all the interconnected submaps. Nodes are overlaid with multiple color bars depending on the number of associated time-points (Figure 6). By providing a visual representation of change in the node expression profile over time, this feature helps users in designing new hypotheses on the role of connected nodes regulating a phenotype.

**Figure 6:**
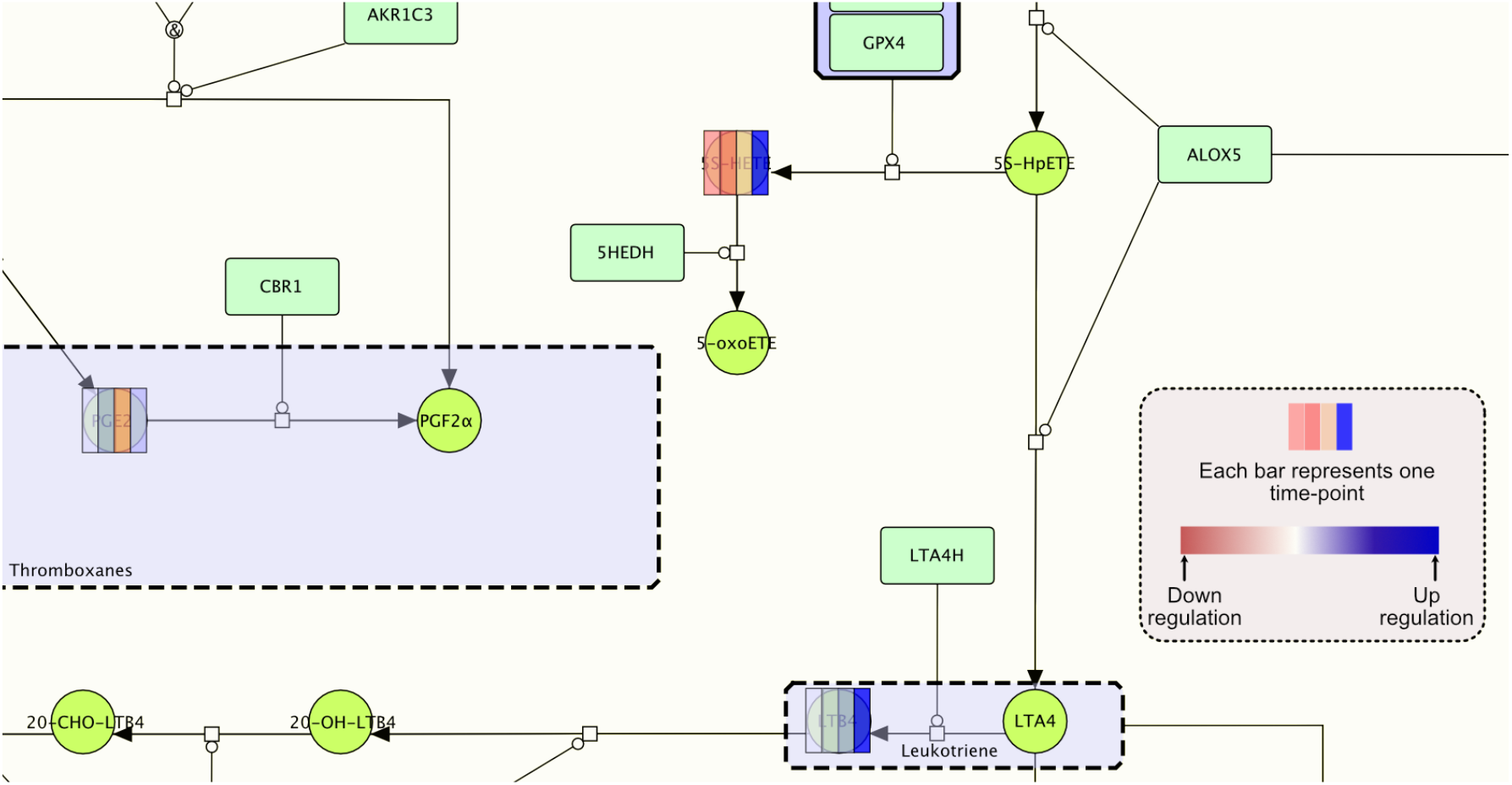
Mapping of time-series data onto the AIR. Nodes are overlaid with colored bar where each bar indicates the data at a given time point. In this example, log2 concentration fold change of selected SPM were calculated at 4 time points (0h, 12h, 24h and 48h) and mapped onto the AIR when mice were challenged with higher titre *E. coli* (10^7^ c.f.u.) compared to self-resolving *E. coli* infection (10^5^ c.f.u.)^43^. The color gradient from red to blue indicates downregulation and upregulation. The color bars demonstrate that all the pro-inflammatory lipid mediators (PGE2, 5S-HETE and LTB4) are mostly upregulated from time point 0h to 48h due to high exposure of *E. coli*.

#### 3.4.4 The identification of core regulatory processes

In the last decade, several methodologies have been developed to define and identify core regulatory network, disease modules, context-specific subnetworks from large molecular interaction network^44–47^. The AIR allows interfacing with such approaches in general through additional plugins. As an example, we provide the user with an interface to identify a core regulatory network from the AIR responsible for the overall dynamics associated with the acute inflammatory process or phenotype under investigation. The detailed methodology for the prediction of the core regulatory network is summarized in our previous publications^23,44^. The methodology is based on the prioritization of feedback loops derived from user-provided multi-omics data, integration of prioritized motifs and finally the preparation of ready to use network file in standardized SBML format for *in silico* analysis.

#### 3.4.5 The AIR in the approximation of aggregate influence on molecular processes / clinical phenotypes due to the change in the level of MIM components

The AIR contains a large number of complexes and placeholders (gene / protein classes known to perform qualitatively similar biological function) regulating biological processes and phenotypes. These processes / phenotypes are described in the AIR from gene ontology databases e.g. Gene Ontology Resource available (http://geneontology.org), Mammalian Phenotype Ontology (http://informatics.jax.org). To facilitate a rapid assessment of the influence of change in the concentration/expression profile of regulatory components on the phenotype level (e.g. increased acute inflammation; increased vasodilation; decreased neutrophil numbers; efferocytosis etc.) at various time-points and/or in various experimental conditions, the AIR implements various logic-based rules to first assign expression profiles to these complexes and place holders. All the submaps present in the AIR are accessible as standard SBML files, which can be analyzed using several state-of-art tools and algorithms to predict the influence on the phenotype. Example of an algorithm that gives a measure of how changes in the levels of components in the MIM contribute to a phenotype is provided in Figure 7. The influence of the MIM components on the phenotype levels may help clinicians in making quick decision on disease state after mapping of patient specific multi-omics data onto the AIR.

**Figure 7:**
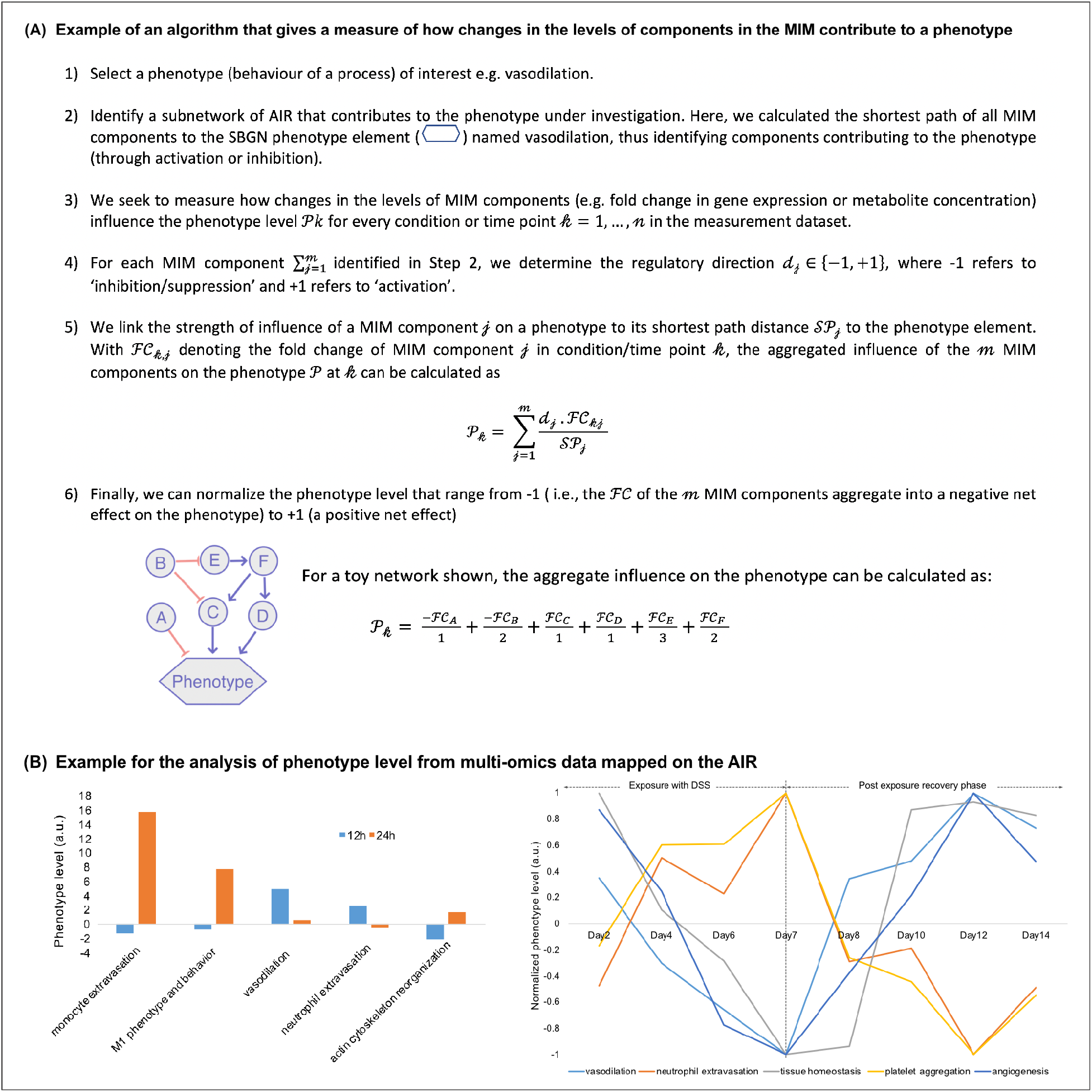
Examples using the AIR for bioinformatic analyses. (A) Algorithm to determine the aggregated influence of change in the MIM components on phenotype level. A toy network in the bottom highlights the aggregate influence on the phenotype due to the expression or concentration fold change in the network components A to F. (B) Examples for the analysis of phenotype level from multi-omics data mapped on the AIR. Left: An example where we measure the influence on the phenotype level (a.u.) from MIM components after mapping of miRNA log fold change data from the zymosan-induced peritonitis mouse model treated with/without RvD1 at time point 12h and 24h^48^. The bar in the plot indicates influence on the phenotype levels when RvD1 was co-administered. The graph indicates that vasodilation was quickly downregulated which is also supported by the low level of neutrophil extravasation. Other phenotypes (monocyte extravasation; M1 phenotype and behavior; acting cytoskeleton reorganization) were upregulated, suggesting that RvD1 brought the whole systems quickly towards the inflammation resolution phase in comparison to the exposure with zymosan alone. Right: In another example, we highlight the influence of MIM components on various processes associated with the acute inflammation onset and resolution after mapping of time-series transcriptomics profile from mouse colitis model^49^. The graph indicates normalized phenotype levels (a.u.) (‘vasodilation’, ‘neutrophil extravasation’, ‘tissue homeostasis’, ‘platelet aggregation’, and ‘angiogenesis’) from mouse model exposed with dextran sodium sulfate (DSS) for 7 days to induce acute colitis followed by 7 days of recovery phase. In this study transcriptomics profiling of colon samples were carried out for 9 different timepoints (Day 0, Day 2, Day 4, Day 6, Day 7, Day 8, Day 10, Day 12 and Day 14). Results suggest that the ‘neutrophil extravasation’ and ‘platelet aggregation’ increases until the DSS exposure (i.e. 7days, inflammation initiation phase) followed by sharp decline during the post exposure recovery phase. On the other hand, ‘vasodilation’ increases from day 7 to 12 (inflammation transition phase) and then a sharp decline in the phenotype was observed suggesting that the system is in inflammation resolution phase.

#### 3.4.6 AIR as a platform to connect inflammation communities

The AIR is freely available to the community on the MINERVA platform hosted on ELIXIR, an intergovernmental organization that brings together life science resources from across Europe (https://air.elixir-luxembourg.org). The AIR also provides an interface where communities can directly raise questions, suggest the inclusion of molecules and processes and update information.

## 4. Conclusions

We have developed a comprehensive *Atlas of Acute Inflammation and Resolution or AIR*, covering more than 30 highly interconnected submaps at the molecular level. Key points summarizing the AIR are provided in Text Box 1.

### Text Box1: The AIR at a glance

- The AIR is the first comprehensive collection of molecular interaction maps underlying acute inflammation initiation, transition, resolution, repair and return to homeostasis.
- The AIR for acute inflammation and inflammation resolution is prepared by extending disease genes associated with primary clinical indications of acute inflammation and known DAMPs, PAMPs with experimentally validated interacting partners.
- The AIR provides biosynthesis and down-stream signaling cascades of protein mediators (e.g. annexin-2, IL-10, TGF-β, INF-α) and lipid mediators (e.g. Prostaglandins, Leukotrienes, Lipoxins, Resolvins, Protectins and Maresins) along with time-series LC-MS-MS data from selected acute-inflammatory phenotypes which can be used for designing new therapeutics.
- The AIR is a portal to other databases (e.g. miRTarbase, UniProt, GenBank, PubMed etc).
- Users can type in the name of protein / gene / regulatory molecule / lipid / biological processes to find the associated functional modules.
- Connections to known drugs and chemicals can be searched directly from the AIR with linked databases.
- Molecular interaction maps available on the AIR are both human and machine readable in a standardized SBGN, SBML format to allow the reproducibility.
- The AIR can be visualized with several regulatory layers including transcription factors, miRNA, lncRNA, drug candidates.
- The AIR itself becomes a knowledge-base to generate hypotheses around acute inflammation and inflammation resolution.
- The AIR provides a scaffold to understand/hypothesize the drug mode of action in inflammation resolution.
- All the edges present in AIR are annotated with PubMed IDs. Thus, all the information present in the AIR is reliable and transparent.
- With visualization of levels of molecular, processes and cellular phenotypes level visualization, the AIR can be used as a tool to translate results from experimental animals to the clinical settings.
- The AIR is enriched with recurring structural patterns called network motifs including feedback/feedforward loops, which induce non-linear dynamics.
- The AIR provides an interface for the inflammation research community to interact.

The AIR is designed with the aim to connect clinicians, biochemists, systems biologists, computer scientists and drug developers. Depending on the need of the end-user, the AIR provides various levels of representation and organization of data and models. Resources such as AIR need continuous improvement with the inclusion of missing links at the molecular level as soon as they are published. This can be realized only through community efforts. The AIR provides an interactive platform to connect the community; thus, we hope that the AIR will be sustained in future by the community.

## Authors contributions

The C.Serhan, S.Gupta and O.Wolkenhauer conceived the manuscript and drafted the first versions. All authors contributed to the scientific content and helped writing the text. All authors approved of the submitted version. O.Wolkenhauer and S.Gupta supervised projects that included the curation of content to the MIM, or layouting submaps. S.Gupta, S.S.Gupta, P.Schopohl, M.Hoch, D.Brauer, F.M.Khan and D.Gjorgevikj designed various submaps. O.Wolkenhauer, S.Gupta, S.S.Gupta and C.N.Serhan equally contributed to the quality check of content. Furthermore, all authors contributed to the interpretation and quality control of information contained in the AIR.

## Data availability

The AIR is hosted on ELIXIR, an intergovernmental organization that brings together life science resources from across Europe and can be accessed through https://air.elixir-luxembourg.org. All the submaps included in the AIR can be directly downloaded in SBML notations. Tutorials to use the AIR are available on https://air.bio.informatik.uni-rostock.de.

## Acknowledgements

The AIR uses Minerva, developed by the Luxembourg Centre for Systems Biomedicine (LCSB). We are grateful for the Minerva support provided by Piotr Gawron and Marek Ostaszeweski. The LCSB is also the elixir node hosting the AIR. We acknowledge Tom Gebhardt and Martin Scharm for providing IT support for the implementation and installation of the AIR. We acknowledge Patrick Schopohl, David Brauer, Krishna Pal Singh, Juliane Procksch, Faiz M Khan, Dragana Gjorgevikj, Julia Scheel and Moritz Kunzmann for their contribution in the curation of content, visualizing submaps and giving directions to interactions from literature. V.Chiurchiù was supported by grants FISM 2017/R/08 and GR-2016-02362380. S.Gupta and O.Wolkenhauer acknowledge support from Bundesministerium für Bildung und Forschung (BMBF) grants [MelAutim (01ZX1905B) and SASkit (012X1903B)] and funding received from the European Union’s Horizon 2020 research and innovation programme under the Marie Skłodowska-Curie grant agreement No 765274. C.N.Serhan acknowledges support from USA NIH GM038765.

## Declaration

The project was in part supported by Heel GmbH. The funders had no role in study design, data collection, curation of content, analysis. The AIR is built from experimentally validated information from the literature, with no information related to products of pharma companies being referred to. All authors declare that there are no competing financial interests, that could undermine the objectivity, integrity and value of a publication.

## Abbreviations

AIR: Atlas of inflammation resolution
DC: Dendritic cell
ILS: Innate lymphoid cells
Mres: Residual macrophage
M1: M1 macrophage
M2: M2 macrophage
Treg cell: Regulatory T cells
PIM: Pro-inflammatory mediators
SPM: Specialized pro-resolving lipid mediators
RvD1: Resolvin D1
PMN: Polymorphonuclear leukocyte
PAMPs: Pathogen associated molecular patterns
DAMPs: Damage associated molecular patterns
SBML: Systems biology markup language
MIM: Molecular interaction map

## Notes

#### Summary of Updates

Typos, choosing an alternative name for one molecule and correction of the names in the list of authors.

https://air.bio.informatik.uni-rostock.de

